# A Fit for Purpose Approach to Evaluate Detection of Amino Acid Substitutions in Shotgun Proteomics

**DOI:** 10.1101/2023.08.09.552645

**Authors:** Taylor J. Lundgren, Patricia L. Clark, Matthew M. Champion

**Author notes:** To whom correspondence may be addressed: Matthew M. Champion or Patricia L. Clark., Department of Chemistry and Biochemistry, 100 McCourtney Hall 140B, Notre Dame, IN 46556, **Email:** or. **Author Contributions:** M.M.C., P.L.C., and T.J.L. designed research; T.J.L. performed research, M.M.C. and T.J.L. performed data analysis, M.M.C., P.L.C., and T.J.L. performed data interpretation, M.M.C. and T.J.L contributed new analytical tools T.J.L., M.M.C., and P.L.C. designed the figures and wrote the paper. **Competing Interest Statement:** The authors declare no competing interest.

## Abstract

Amino acid substitutions (AAS) change a protein from its genome-expected sequence. Accumulation of substitutions in proteins underlie numerous diseases and antibiotic mechanisms. Accurate global detection of substitutions and their frequencies would help characterize these mechanisms. Measurement of AAS using shotgun proteomics is attractive due to its high sensitivity and untargeted acquisition. However, identifying substituted peptide-spectra requires search strategies that extrapolate beyond the genome, which can introduce bias. To characterize this bias, we constructed a “ground-truth” approach using the similarities between the *Escherichia coli* and *Salmonella typhimurium* proteomes to effectively model the complexity of distinguishing substitutions from genomic peptides. Shotgun proteomics on combined whole cell lysates from both organisms generated a library representing nearly 100,000 peptide-spectra and 4,161 distinct peptide sequences corresponding to genome-level single AAS with defined stoichiometry. We tested the ability to identify *S. typhimurium* peptide-spectra using only the *E. coli* genome in substitution-tolerant database searching. Overall, 64.1% of library peptides were correctly identified. We observed a wide range of identification efficiencies based on the specific AAS, but no inherent bias from stoichiometry of the substitution. Short peptides and substitutions near peptide termini, which require specific diagnostic ions for unambiguous identification, are matched with below-average frequency. We also identified “scissor substitutions” that gain or lose protease cleavage sites. Although scissor substitutions are chemically distinct from the genomic peptide, they had poor identification efficiency. This ground-truth AAS library identifies multiple sources of bias in AAS peptide-spectra identification and sets expectations for the application of shotgun proteomics to testing AAS hypotheses.

**Significance statement:** High-fidelity decoding of the genome is essential for life. Mistranslation leads to amino acid substitutions, which can disrupt protein folding and function, and impact cell fitness. Detection of mistranslated protein products necessitates robust and non-biased approaches. Proteomics is a promising solution, but identifying non-genomic peptide-spectra is a severe bioinformatics challenge. We created a ground-truth library of substituted amino acid peptides by mixing two closely related bacteria in a single sample. We quantitatively defined the degree to which informatics could correctly distinguish substituted peptides when single-organism databases are present. This approach defines intrinsic and informatics limits in substitution detection in shotgun proteomics and identifies previously overlooked challenges with identifying “scissor substitutions”.

## Introduction

Incorporation of an incorrect amino acid during translation can negatively impact protein stability and function, leading to increased misfolding and proteotoxic stress.(1) Several oncogenes increase mistranslation through modification of mRNA and tRNA.(2–5) Similarly, several antibiotics create mistranslation-driven proteotoxic stress, which compromises the cellular membrane of bacteria.(6) However, in both cases the distribution and sequence biases of the mistranslated proteins are largely unknown, due to the difficulty of identifying diverse, rare substitutions in the background of a genome-defined proteome. This uncertainty restricts informed improvement of therapeutics that target translational fidelity. Defining proteotoxic products of unfaithful translation requires untargeted discovery of nongenomic modified proteins. Global detection of amino acid substitutions in proteins is of particular interest to evaluate translation fidelity and the consequences of mis-translation on protein homeostasis.(1, 7–9)

Shotgun proteomics is a powerful technique for untargeted protein identification.(10, 11) However, peptide-spectra are predominately identified by matching to a genome-defined database and subsequent protein inference.(12) Identification and inference beyond genome-anticipated peptides introduces a significant bioinformatic challenge, which complicates amino acid substitution (AAS) identification. Classical permutation-based searching for stochastically modified amino acid residues in peptide-spectra is only effective when few modifications are considered.(13, 14) The full suite of single AAS permutations dramatically expands the search space to be intractable to permutation-based search. For example, an average tryptic peptide of 14 aa times 18 mass-unique substitutions equals 252 possible canonical single AAS permutations, before considering other common post-translational peptide modifications that may also occur. Alternatively, *de novo* peptide sequencing is not constrained by the genome but this approach lacks the identification power of a database-driven search.(15–17)

Other proteomic search approaches have successfully identified substituted peptides. Previously, many hundreds of substitutions were identified using targeted databases informed by mRNA sequencing or oncological single nucleotide polymorphisms.(18–20) Novel search algorithms have enabled the detection of many peptide modifications, including AAS, using only the reference genome. Recently, Mordret *et al*. adapted dependent-peptide search to identify 1,679 unique substitutions.(7) There are also reports successfully identifying substitutions from reference genomes using proprietary commercial software, such as the SPIDER algorithm in PEAKS.(6, 17, 21, 22)

The current paradigm for evaluating software identification of peptides is limited to the range and yield of peptide-spectra matches (PSMs) at controlled false discovery rates (FDR).(23) Yet the yield of detected AAS PSMs is an information-poor metric, because it cannot characterize nor count un-or mis-identified AAS peptide-spectra. A comprehensive description of un- and mis-identified spectra can only be accomplished by *a priori* knowledge in a ground-truth dataset, which is necessary to describe the limitations introduced by identification of AAS-peptide spectra. This description could also evaluate logical assumptions made about AAS identification such as inferred lower limits of detection, the impact of peptide stoichiometry, and the integrity of FDR estimation.

Previously used positive controls for AAS detection include incorporating synthetic peptides and customized search databases.(6, 19, 21) These have been effective for targeted validation of successfully identified AAS PSM, however both are limited in the number of representative AAS peptide sequences that can be reasonably included in an experiment due to price and expansion of the search space, respectively. (24) This makes it challenging to represent each substitution type across diverse peptide or protein localization and across unique competition from sample matrix, other peptides, and artificial modifications. To reflect the majority of AAS, a positive control for global identification of AAS would be diverse in substitution type, location within the peptide, absolute and relative abundance.

To recreate the broad range of substitution and peptide contexts, we leveraged naturally occurring diversity to create a useful positive control for global AAS detection. Many proteins have homologs in closely related species, with naturally occurring amino acid polymorphisms between species defined by their respective genomes.(25) We hypothesized that combining two closely related species would result in a subset of anticipated AAS peptides of sufficient complexity and diversity to represent the physiochemical characteristics observable in a shotgun proteomics experiment (Figure 1A). Peptides from both organisms could be identified in a single standard database search where both genomes are provided, resulting in a library of positively identified spectra that represent a substitution (Figure 1B). The same data could then be searched with only one genome and an AAS peptide-spectrum identification strategy of choice (Figure 1C). Pairwise comparison of searches by spectrum allows evaluation of the AAS search strategy by global and categorical efficiency at controlled stoichiometry.

**Figure 1.**
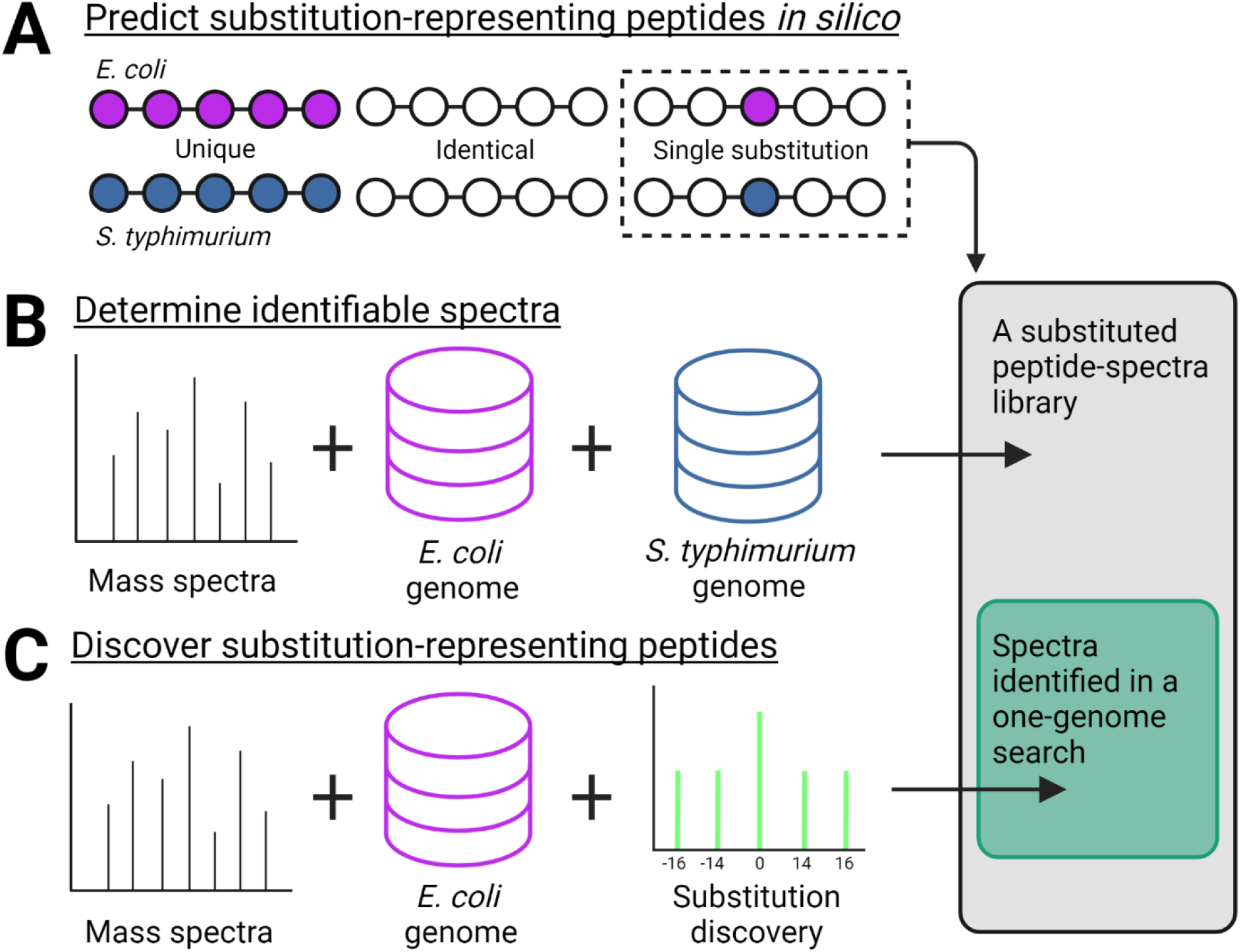
Creating the amino acid substitution (AAS) peptide-spectra library. **A)** Peptide sequences that mimic an AAS are predicted from *in silico* digested *E. coli* and *S. typhimurium* protein sequences. **B)** Peptide-spectra from a mixed proteome sample are identified by a standard database search with both genomes provided. Identified spectra that match the sequence of a single AAS peptide are used to create a peptide-spectra library, used as a positive control for AAS discovery search. **C)** The same spectra are searched with only one genome and a method to discover AAS peptides. We used MSFragger’s mass-offset search to identify *S. typhimurium* peptides when provided only an *E. coli* genome, or vice versa, and evaluated identification performance against the ground-truth library.

We selected *Escherichia coli* and *Salmonella typhimurium* for this approach, as these bacteria have complete genome sequences and sufficient evolutionary distance for unambiguous protein inference.(26, 27) This resulted in a ground-truth library of 52,756 AAS representing spectra from *Salmonella* that we used to discover biases in AAS identification. We adapted the mass-offset approach to identify AAS and found moderate success, with 64.1% peptide identification efficiency.(28) We found successful PSMs had similar absolute and relative abundance sensitivity but scored lower than their library counterparts. Furthermore, we identified both substitution type and location as driving biases in PSM identification efficiency. We also define “scissor substitutions”, a unique subclass of substitutions that remove or introduce a protease cleavage site, which resulted in fewer successful identifications. We demonstrate the utility of a spectral library and the power of efficiency as a criterion for evaluating limitations of AAS identification.

## Results

### Constructing the ground-truth substitution library

To evaluate the identification of substitutions in shotgun proteomics, we needed a set of peptide-spectra that met three criteria. First, each spectrum needs sufficient evidence to be confidently assigned an amino acid sequence. Second, each spectrum must represent a peptide sequence that differs by one amino acid from a reference genome. Third, the collection of spectra should represent substitutions diverse in substitution type, location, and sequence context. We found that by mixing two closely related bacteria, *E. coli* and *S. typhimurium*, evolutionary homology would result in a ground-truth library that broadly satisfies these three constraints. To identify genome-defined peptides that vary by a single amino acid between organisms, both genomes were digested to proteotypic peptides *in silico* using Protease Guru, then filtered to exclude exact peptide homologs and peptide sequences that differ by more than one amino acid (see Materials & Methods).(29) We found 31,053 *S. typhimurium* peptides that represent a single amino acid variant (SAAV) against an *E. coli* cognate sequence and 30,756 *E. coli* peptides that represent a SAAV against a *S. typhimurium* cognate sequence (Figure 1A). However, we expect that many of these substitution-representing peptides will not be observable under a single experimental condition for reasons including low gene expression, incomplete digestion enzyme efficiency, and peptide characteristics incompatible with efficient ionization.(30, 31) To collect spectra representing these peptide sequences we prepared, quantified, and digested *E. coli* and *S. typhimurium* whole cell lysates individually using standard shotgun proteomic preparation methods. Digested lysates were combined and serially diluted, measured via nUHPLC-MS-MS/MS, and a standard database search performed with both the *E. coli* and *S. typhimurium* genome. This resulted in 52,756 spectra and 2,568 unique peptide sequences that comprise our ground-truth library (Figure 2A). This library shows considerable diversity in physiochemical properties (Supplemental Figure 1) and represents 241 of the 342 possible amino acid substitutions detectable by mass spectrometry (Supplemental Figure 2). This ground-truth library meets our three criteria and is representative of AAS observable in typical shotgun proteomic experiments.

**Figure 2.**
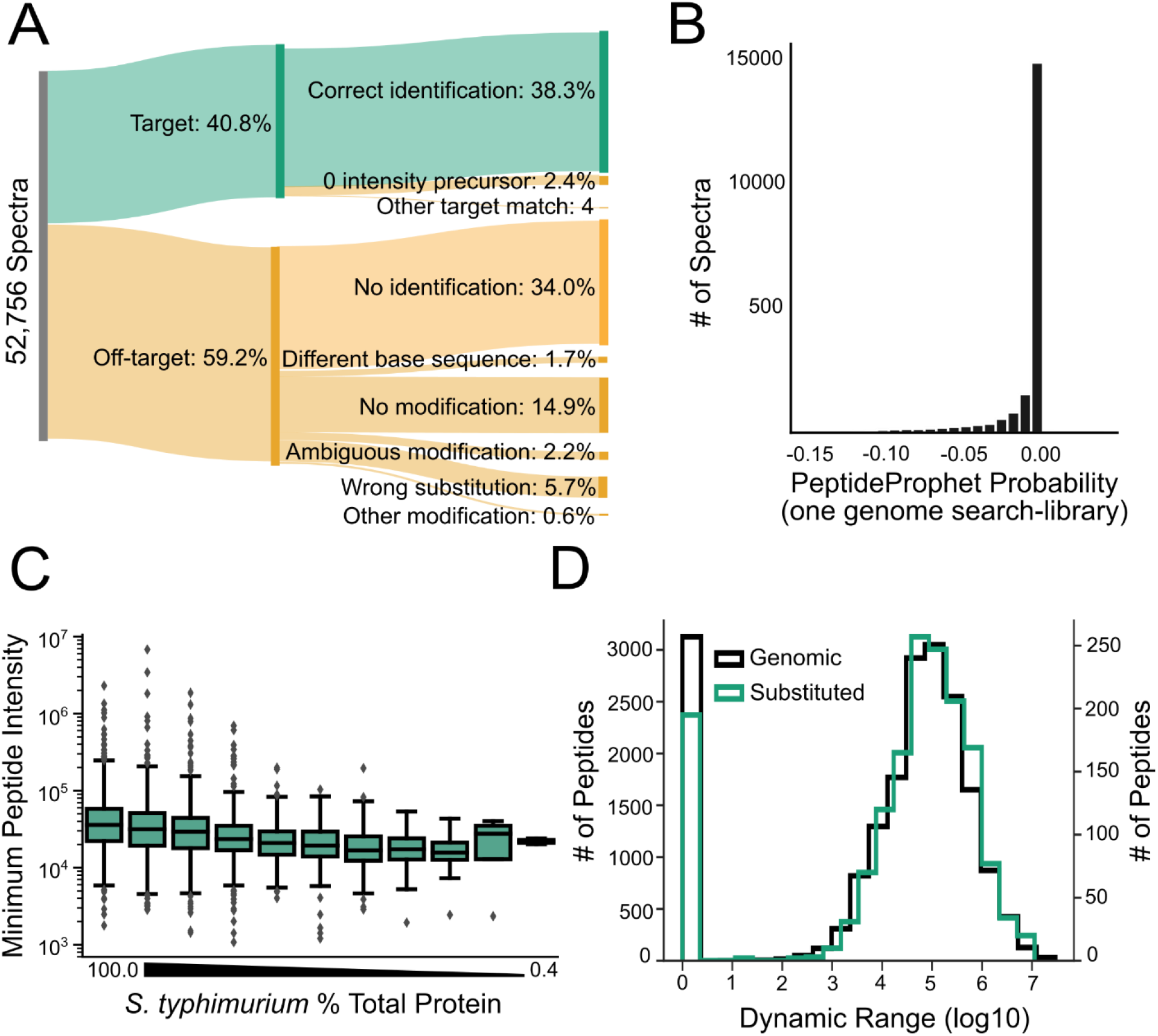
Tracing the final fate of library spectra in the one genome search demonstrates intensity independent challenges for spectrum identification. **A)** The final fate of each library spectrum identification in the one genome search. Spectra were first categorized as representing a library peptide sequence (green) or other sequences (gold). Spectra were further categorized as a correct identification (green), or an incorrect identification (gold). The sub-categories of incorrectly identified spectra represent either failure to identify the spectra or specific competing identification hypotheses in sequence assignment. These categories can be used to guide search software improvements. **B)** The PeptideProphet score difference (one genome search minus library) for each correctly identified spectrum. There is a tail of spectra that scored more poorly in the one genome search. **C)** The minimum observed intensity for each peptide, grouped by the final sample of the dilution series that led to successful identification. The distributions of minimum observed peptide intensities were similar throughout the dilution series, despite decreasing *Salmonella* stoi-chiometry. **D)** The dynamic range observed for each correctly identified AAS peptide (hatched white bars) and all *S. typhimurium* peptides (black bars) though the dilution series. Both peptide groups have a similar observed dynamic range distribution.

### Determining the efficiency of mass-offset discovery of amino acid substituted spectra

We applied our ground-truth substitution library to determine the efficiency of AAS identification in shotgun proteomics. We used mass-offset search to identify *S. typhimurium* peptide-spectra using only an *E. coli* search database (Figure 1C). Similar to dependent peptide search in Mordret et al.(7), we adapted mass-offset PTM search functionality in MSFragger to identify AAS peptides (see Materials & Methods).(11) We created a Python script (Materials and Methods) to annotate identified mass-offsets with specific change in mass and aa localization as substitutions while filtering out mass-ambiguous modifications and unquantified peptides. We tracked the fate of each individual library spectrum in the single-genome search and evaluated identification efficiency globally and categorically. Most library peptide sequences (64.1%) were identified, though only 38.3% of library spectra were correctly identified (Figure 2A, in green). Many spectra (34.0%, ‘No Identification’) did not score well enough for sequence assignment after standard FDR control. This demonstrates a challenge to obtain confidence in sequence assignment without *a priori* genomic knowledge. The remaining spectra (27.7%, in gold) were confidently assigned an incorrect sequence, whether matched to the unmodified genomic cognate sequence (14.9%), assigned an incorrectly modified sequence (5.7%), or filtered out due to mass ambiguity with other PTMs (2.2%) or lack of intensity (2.4%). These categories represent specific targets for iterative improvement of the identification software and are broadly applicable to any search software. The fates of library spectra suggest that most spectra are un-or mis-identified because spectral evidence of a substitution generates less confidence in spectrum identity than alignment with the genomic database.

We investigated if simply enumerating the mass-offset was sufficient to identify each substitution type. The one genome search did not identify any spectra of 45 substitution types, though most (32 substitutions) had a low number of sample spectra (n<50) in our library (Figure 3A, Supplemental Figure 2). For example, no spectra representing a substitution from Cys were successfully identified. Substitution types with no PSMs in the one genome search but many in the library include substitutions that are isobaric with common modifications, such as T→A (1,000 PSMs), D→ Q (621 PSMs) and Q→ D (529 PSMs). Identified substitution types had a broad range of efficiency from 3.6% (R→ I/L) to 100% (H→ M) (Figure 3A). Likewise, many library spectra representing a substitution involving Arg or Lys had below-average identification efficiency in the one genome search. We conclude that mass-offset identification is generally suitable for global substitution identification, but that special care should be taken for application to identification of specific substitution types.

**Figure 3.**
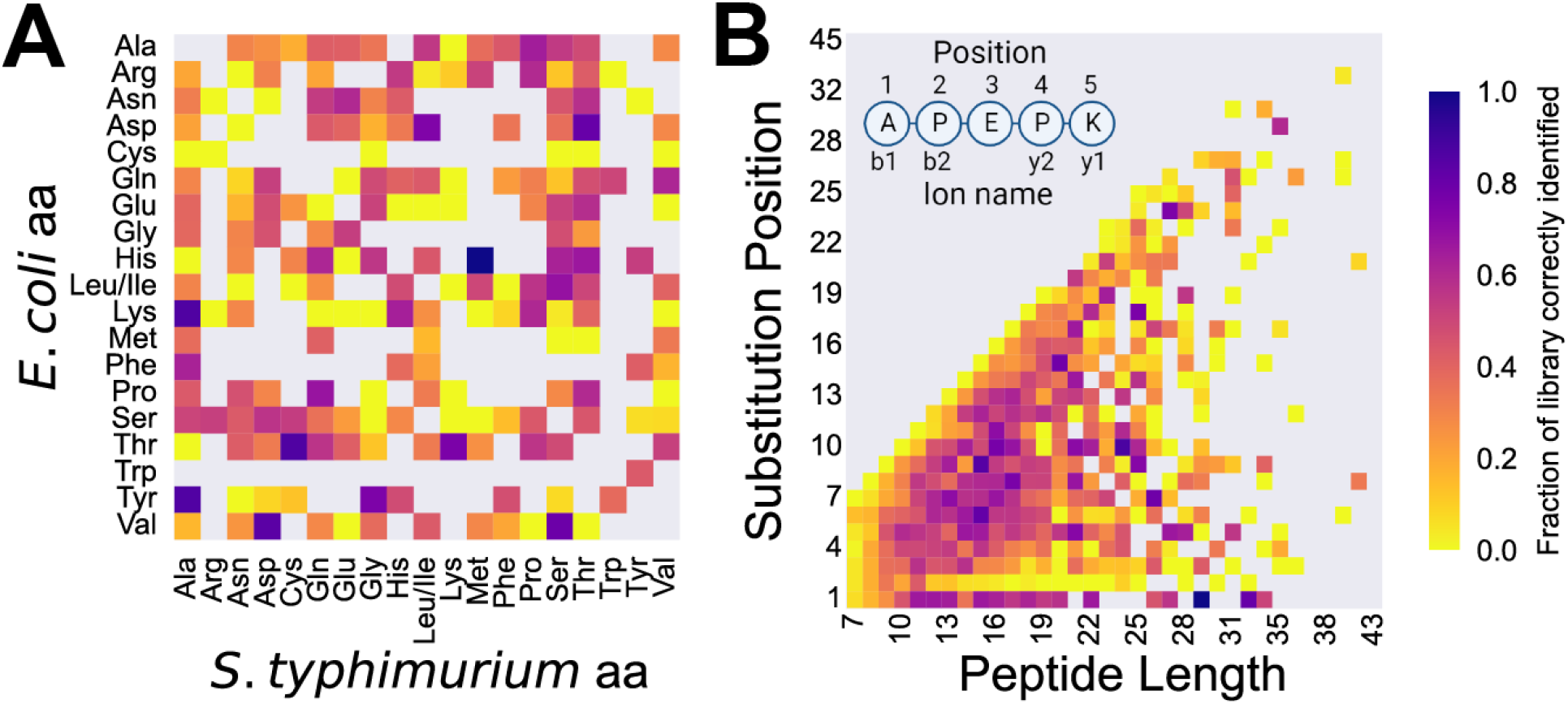
Efficiency of library spectra identification demonstrates categorical bias in AAS identification. **A)** AAS library spectra identification efficiency by substitution type, comparing the genome-anticipated *E. coli* aa to the aa present in the *S. typhimurium* peptide. Substitution types absent in the library (grey) include aa highly conserved in bacteria such as Cys, Phe, and Trp. Substitutions involving Arg, Cys, or Lys had below average identification efficiency. **B)** AAS library spectra identification efficiency by substitution position and peptide length. Substitution position and peptide length combinations not represented in our library are shown in grey. Substitution position 1 represents the N-terminal aa; the y=x position represents the C-terminal aa. Substitutions in the middle of moderately sized peptides had the highest identification efficiency. In contrast, small peptides (<9 aa), large peptides (>25 aa), and substitutions at the b_2_, y_1_ or y_2_ positions had poor identification efficiency.

### Successfully identified spectra met higher score thresholds with lower individual scores

We hypothesized that the inability to identify many spectra in the single genome search was due to an increased burden of proof without *a priori* sequence knowledge. Spectra sequence assignment uniquely depends on a PSM score relative to a cutoff value. The score cutoff thresholds are universally based on the target-decoy strategy.(23) To investigate if identifying substitutions with mass-offset affected the target-decoy resolution, we compared the distribution of decoy PSM scores between the one genome and library search (Supplemental Figure 3). There are more decoys with higher scores in the one genome search, indicating that higher score thresholds are necessary to maintain confidence with a controlled false discovery rate. Next, we asked if the same spectral evidence was uniquely weighted by the mass-offset expansion of search space. To do this, we parsed the library spectra correctly identified in the one genome search and took the difference of library search PeptideProphet Probability score from the one genome search score (Figure 2B).(32) Although the plurality of correctly identified spectra received the same score in both searches, many spectra received a lower score in the single genome search. This indicates mass-offset peptides are disadvantaged in the scoring algorithm. Together, these results demonstrate that AAS peptide-spectra score more poorly and require higher thresholds for successful sequence identification via mass-offset.

We next sought to determine the driving factors that distinguish neutral and score-disadvantaged substitution identifications. We first suspected peptide abundance, as a secondary factor of scoring is MS2 fragment ion signal intensity. Co-eluting peptides provide competitive ions that may mask identification of low abundance peptides. To determine if AAS identification efficiency was uniquely affected by stoichiometry, we diluted *S. typhimurium* lysate two-fold against a constant background of *E. coli* lysate (Figure 2C, D). We calculated the dynamic range for all *S. typhimurium* peptides as the log difference between maximum and minimum observed precursor intensity per peptide sequence across all dilutions. We observed a similar dynamic range between *S. typhimurium* AAS representing peptides discovered in the single-genome search and all *S. typhimurium* peptides identified in the library search (Figure 2D). We next asked if substitutions had a unique lower limit of identification, or a stoichiometry-dependent limit of identification. We determined the lower limit of abundance for peptide identification by plotting the intensity distribution of peptides at their last identification in the dilution series (Figure 2C). The minimum detected *S. typhimurium* peptide intensity distribution was similar across decreasing stoichiometry. The similar dynamic range and lower limit of identification imply that identification of substituted peptide-spectra is limited by conditions that reduce identification of all spectra and is not uniquely disadvantaged by abundance nor stoichiometry.

### Specific fragment ions are required for confident assignment of some substituted peptide sequences

We next looked for unique characteristics of the library spectra not correctly identified in the single genome search. The distributions of peptide intensity, retention time, ion mobility, shift in retention or shift in ion mobility from the cognate peptide were similar between successfully identified library spectra (Supplemental Figure 1) and incorrectly or unidentified library spectra (Supplemental Figure 4). The remaining characteristics of a spectrum, fragment ions and their intensities, are uniquely weighted during AAS discovery. Lacking *a priori* sequence knowledge imposes a burden of proof on specific fragment ions diagnostic of peptide modification. We identified two scenarios where this burden of proof decreased AAS identification efficiency: substitutions near the peptide terminus, and ambiguous localization of isobaric mass-offsets.

**Figure 4.**
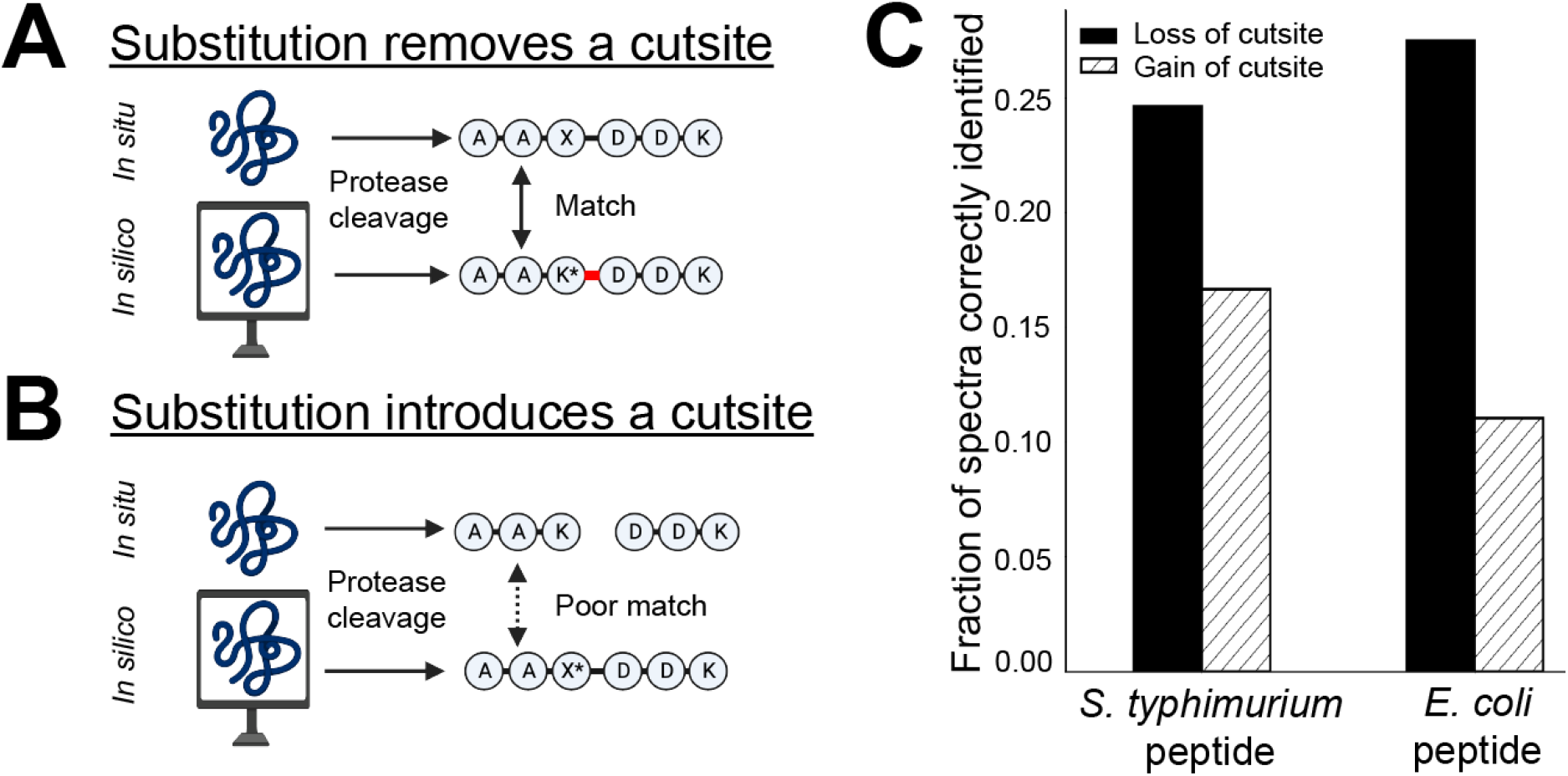
Identification of scissor substitution peptide-spectra is disadvantaged by peptide search engine logic. Scissor substitutions, which add or remove a cutsite for the protease used to generate peptides, are disadvantaged by singular digestion of proteins *in silico*. **A)** Scissor substitutions that remove a protease cleavage site result in a physical peptide that matches an *in silico* missed cleavage peptide (in red), a scenario commonly considered in search software. **B**) In contrast, scissor substitutions that add a cut site result in a physically cleaved peptide. Peptide prediction from protein sequences typically occurs before annotation of identified substitutions, resulting in an undigested *in silico* peptide and no detection of the expected physical peptides. **C)** The identification efficiency of AAS PSMs for each class of scissor substitution. Very few scissor substitution-representing *S. typhimurium* peptides (left) that gain a new tryptic site (white hatched bars) were identified in our search using only the *E. coli* genome. As expected, this pattern was reciprocated when *E. coli* peptides representing a scissor substitution (right) were identified using only the *S. typhimurium* genome, implicating the peptide identification search logic.

Each amino acid in a peptide contributes to the mass of multiple fragment ions. For example, the N-terminal residue is part of the mass balance of every b ion generated in MS2 but does not contribute to any y ions. AAS near either terminus would generate few complimentary, modified b/y ion pairs that provide strong evidence for sequence assignment. Additionally, the number of ions potentially diagnostic for a substitution increases with peptide length. We hypothesized that these constraints would introduce position and length biases on single genome search AAS peptide identification. To determine if substitution location influenced the identification of AAS spectra, we plotted the efficiency of library spectra identification by length and substitution position (Figure 3B, number of representative spectra presented in Supplemental Figure 5). We found short (<9 aa) or long (>23 aa) peptides; also, substitutions at the b_2_, y_1_ or y_2_ positions were identified at below average efficiency. N-terminal substitutions were robustly identified, comparative to the average success rate. This may be caused by the software’s arbitrary assignment of substitution position to the aa closest to the N-terminus when the precise residue cannot be determined. The C-terminal substitutions we identified were limited to the swap of lysine and arginine. Other C-terminal peptide substitutions would lack a protease cleavage site. Substitutions at these positions in the protein sequence were identified in the longer peptide with a missed protease cleavage and a substitution at the missed cleavage residue (see section on scissor substitutions, below).

Isobaric substitutions also require specific diagnostic ions for proper localization and identification. For instance, a G→ A substitution may be confused with a S→ T, D→ E, N→ Q, or V→ I/L substitution, as each represents a mass-offset of 14.016 Da. If both G and S are in the peptide sequence, unambiguous identification requires intensity from the y or b ions between these residues (Supplemental Figure 6A). These ambiguous localizations of isobaric mass-offsets account for many of the 3,054 spectra assigned an incorrectly substituted sequence. To better understand the mass degeneracy within AASs, we plotted the number of substitution mass shifts within ±0.02 Da for each substitution type (Supplemental Figure 6B). We found 190 out of 342 substitution types were isobaric with at least one other substitution, while some had up to five unique substitutions representing the same change in mass. Surprisingly, many multiply degenerate substitutions had average or better library spectra identification efficiency (compare Figure 3A, Supplemental Figure 6B), likely due to strong localization of the mass-offset. The remaining misidentified spectra suggest the application of other dimensions of data, such as retention time or ion mobility, to disentangle isobaric modifications.

### Scissor substitutions conflict with how peptides are predicted from protein sequences

Substitutions that add or remove a protease cleavage site, such as a substitution to or from lysine in a tryptic digest, result in peptides of different length and sequence than their genomic cognates. We call these “scissor substitutions” and hypothesized that their detection is disadvantaged by how search database peptides are predicted from proteins. Genome-defined protease cleavage motifs are identified *before* identifying spectra, and peptide modifications are determined without reconsideration of peptide cleavage. For scissor substitutions that lead to loss of a cutsite, spectrum identification requires the consideration of a missed cleavage *in silico* for an anticipated, modified peptide to match the physical peptide (Figure 4A). An example annotated PSM and extracted ion current for this substitution type is provided (Supplemental Figure 7). In contrast, scissor substitutions that introduce a new cutsite cause a discrepancy between the shorter cleaved peptides *in situ* and the longer *in silico* anticipated sequence, with no additional consideration currently available to adjust software expectations (Figure 4B). In agreement with this logic, we found the library spectra identification efficiency for spectra representing the gain or loss of a cutsite was below average (Figure 4C). There is a marked distinction between identification of substitutions resulting in the loss of a cutsite (24.6%) and the gain of a cutsite (16.7%, Figure 4C). To ensure poor identification of library spectra representing the gain of a protease cleavage motif was independent of the representative *S. typhimurium* peptide contexts, we performed a reciprocal search to discover *E. coli* PSMs using only the *S. typhimurium* genome. As expected, we again observed below average identification efficiency of substitutions that remove a cutsite (27.5%) and dramatically low efficiency for substitutions that introduce a cutsite (11.1%, Figure 4C).

## Discussion

Applying our mixed organism ground-truth library to evaluate identification of substituted peptide spectra using one genome and the mass-offset search strategy revealed the global and categorical efficiency of software analysis, which should inform both software and experimental adjustments for improved sensitivity. In our work, we set an initial benchmark of 64% substituted sequence coverage for a complex and unfractionated data set. Achieving increased proteome depth via in-solution or gas phase fractionation is likely to improve library peptide identification efficiency. Our library of substituted peptides behaved like other peptides, with no unique limits of abundance and stoichiometry on their identification. The mass spectrometer is agnostic of ion identity when isolating and fragmenting peptides. Furthermore, substituted peptides occupy a distinct analytical space of mass, retention time, and ion mobility from their genomic cognate peptides. Thus, the problem of abundance and matrix competition is not solely cognate driven, but rather based on all proteomic interference. Based on the observed frequency of substitutions in other works, we expect many substitutions to have low abundance and signal intensity relative to the proteome.(7, 21) Therefore, approaches such as fractionation known to improve dynamic range and sensitivity should be employed.(33–36)

From these data, the biggest limitation of AAS identification was PSM score. The modal fates of incorrectly identified library spectra were those not confidently assigned a sequence (34%), followed by spectra with enough evidence for the cognate sequence but not for a modification indicative of a substitution (15%). Selective score boosting of bona fide AAS spectra could be accomplished by application of other dimensions of data. There are existing algorithms for prediction of retention time and ion mobility based on peptide sequence.(37, 38) Likewise, there are scoring models that can account for changes to retention time, such as Percolator.(39) In addition to boosting the score of bona fide AAS spectra, these approaches would also decrease confidence in decoy sequences matching to spectra based on mass alone. Combining these data and existing or novel analysis tools may provide better score-based resolution of true AAS PSMs, other peptide sequence assignment hypothesis, and decoy AAS PSMs.

The categories of peptide-spectra misassignment described here delineate other competing hypothesis of spectra peptide sequence assignment. While not designed to be an exhaustive list, these categories captured all the misassigned spectra in our experiment. The two modal categories, unmodified and incorrect substitution assignment, likely reflect the specific fragment-ion bias found in our negative results. Both the length of peptide and the relative position of the substitution can have a dramatic effect on the number of potential ions that unambiguously identify the aa sequence, as opposed to other ions that support all the similar sequence hypotheses. The additional dimensions of mass spectrometry data may alleviate the additional burden of proof on specific fragment ions. While the observed ion mobility shifts between AAS peptides in our work was small, it has been demonstrated that these shifts are larger near the peptide termini, and may boost the identification efficiency of substitutions at the b_2_ or y_2_ positions.(38) Alternatively, substituted peptide sequence coverage can be improved by splitting a sample and digesting each portion with complimentary proteases. The new set of peptides in the parallel digest will have unique lengths and relative substitution positions, providing additional independent evidence of a substitution.

We found that the protease used to generate peptides also defines a subset of substitutions that we term scissor substitutions, which remove or introduce a protease cleavage motif. Scissor substitutions present a significant order of operations challenge that is easy to identify in hindsight but difficult to address *a priori*. Aligning software expectations to physical peptides that lose a cutsite is already possible in virtually every search engine by *in silico* missed cleavage. Despite this logical alignment, and despite not being located near the peptide termini, these library spectra were still identified with below average efficiency. We do not have a solution to align software expectations for substituted peptides that introduce a cleavage site. The addition of modification annotation and second cleavage would only recover spectra identifications for peptide sequences already identified by other spectra. Some of these are explained by stochastic missed cleavages *in vitro*. It is much easier to compensate for scissor substitutions during peptide preparation, by splitting a sample to be digested by two complimentary proteases. Poor substitution coverage at the cleavage motifs of one enzyme would be bolstered by improved coverage in the other digest.

Our ground-truth positive control provides a template for the evaluation of diverse global identification strategies for amino acid substitutions against the current gold standard of database driven search. This approach demonstrates for the first time fundamental factors that uniquely limit identification of substituted peptides, namely; PSM score, substitution type, and a specific fragment-ion burden. These limitations suggest maximizing peptide sensitivity and splitting a sample for digestion with complimentary proteases for improved substituted peptide sequence coverage. Significant work remains for confidence in missing values, where a targeted approach is advisable. Shotgun proteomics is thus a promising tool for positive hypothesis testing regarding global identification of amino acid substitutions.

## Materials and Methods

### Peptide preparation

Overnight cultures of *Salmonella typhimurium* LT2 cells or *Escherichia coli* MG1655 were diluted in LB media, grown to log phase, and pelleted by centrifugation. Next, the pellet was suspended in lysis buffer and lysed using a bead beater (Biospec). Cell lysate was clarified by centrifugation and quantified using a Pierce BCA assay (Thermo Fisher) per manufacturer’s protocols. Samples were reduced, alkylated, and subsequently loaded onto S-Trap mini columns (ProtiFi) per manufacturer’s protocol. Proteins were digested with 1 μg trypsin in 160 μL of 100 mM TEAB pH 8.5 for 2 h at 47°C. Peptides were eluted per manufacturer’s protocol and dried down to ∼20 μL in a speedvac. *S. typhimurium* peptides were desalted using an Oasis HLB desalting column (Waters) and *E. coli* peptides desalted using C_18_ ZipTip (EMD Millipore) following the respective manufacturer’s protocol. Desalted peptides were dried down in a vacuum concentrator, then suspended in 0.1% formic acid to a final concentration of 300 ng/μL.

### Serial dilution of *S. typhimurium*

Desalted *S. typhimurium* peptides were serially diluted two-fold by addition of 60 μL peptides to 60 μL of 0.1% formic acid. Desalted *E. coli* peptides (30 μL) were added to each *S. typhimurium* dilution to create a constant background of *E. coli* peptides.

### Liquid chromatography-mass spectrometry

Technical duplicate injections of 1.33 μL per sample were separated with a PepSep TEN C_18_ 10 cm x 100 μM column (Bruker) and eluted with a 90 min segmented linear gradient. Mass spectra were collected on a Bruker TIMS-TOF Pro operating with the default DDA-PASEF 1.1s cycle time method with two modifications: the CaptiveSpray source set to 1700V and collision energy maximum to 70 eV.

### Identification of mass spectra in FragPipe

Raw data was searched using FragPipe (v.17.1) GUI with MSFragger (3.4) and filtered with Philosopher (v4.2.2-RC). Software parameters for each search are included in the MassIVE repository (See fragpipe.config file). *E. coli* MG1655 (UP000000625) and *S. typhimurium* LT2 (UP000001014) genomes were downloaded from Uniprot (2022.03.25) with common contaminants and decoy sequences added in FragPipe. The two-genome search included up to one missed cleavage, oxidation of methionine as a variable modification, and no mass-offsets. The single genome search in MSFragger was set to use the mass-offset algorithm with a corresponding offset for each AAS and the top 26 PTMs discovered in a default setting open search. The option to report mass-offset as a variable modification was set to 1 (“Yes – and remove delta mass”).

### Genomic analysis to identify single amino acid variant peptides

To identify a target list of tryptic peptides that differ by one aa between organisms, each genome was digested *in silico* using Protease Guru.(29) The resultant list of peptides was input in a Python script (FindSSP.py) that outputs a target list of all peptide sequences that represent a SAAV between the two organisms. The script excludes the mass ambiguous substitutions I/L→ L/I and R/K→ X!R/K at the peptide C-terminus.

### Annotation and filtering of amino acid substitutions

Spectra representing AAS between *E. coli* and *S. typhimurium* were parsed and filtered using a Python script (MSFraggerFindSubs.py). To summarize, all PSMs were imported from MSFragger’s psm.tsv outputs. Modified sequences were matched from the indicated mass-offset to all considered modifications within 25 ppm peptide mass error. For example, APEPT[-18]IDEK would be annotated as ‘T→ A or Dehydration’. All substituted sequences were then filtered to only include target peptide sequences identified by FindSSP.py. PSMs were filtered to remove unquantified peptides with 0 intensity precursors, ambiguous or non-AAS modifications.

## Code and data availability

Data used to generate figures and python scripts used in this work are available at https://github.com/ChampionLab/substitutionannotation. Raw spectra outputs are available at the MassIVE repository (<u>doi:10.25345/C5416T88R</u>).

## Acknowledgements

We thank Dr. Aleksi Nesvizhskii and Dr. Fengchao Yu for assistance with using FragPipe, Dr. Jim Slauch for the kind gift of *Salmonella*, Daniel Hu for advice, Dr. Patricia Champion for critical reading of the manuscript and Dr. Bill Boggess in the Notre Dame Mass Spectrometry and Proteomics Facility for technical advice and assistance. P.L.C. acknowledges support from NIH award DP1 GM146256. M.M.C. acknowledges support from R01GM139277. T.J.L. acknowledges support from NIH training grant T32GM075762.

## Notes

ftp://massive.ucsd.edu/MSV000092029/

### Competing Interest Statement

The authors have declared no competing interest.

