## Supplemental Figures for "A Fit for Purpose Approach to Evaluate Detection of Amino Acid Substitutions in Shotgun Proteomics"

**This file includes:**

Supporting text
Figures S1 to S7
Table S1
SI References

### Supporting Information Text

### Materials and Methods.

Cell Culture

*S. typhimurium* LT2 cells were cultured overnight in LB media, diluted 1:25 and grown to an OD600 of 0.34. A soft pellet was formed by centrifugation at 5,000 x*g* for 10 minutes at 4°C, and supernatant decanted for a remainder volume of 90 mL. The pellet was resuspended with shaking, split evenly between two 50 mL tubes, then pelleted again by spinning at 7,000 x*g* for 15 minutes at 4°C.
*E. coli* cells were cultured overnight in LB media, diluted and grown to an OD600 of 0.6. They were pelleted by spinning at 7,000 x*g* for 15 minutes at 4°C.

Lysis

Cell pellets were suspended in 1 mL of lysis buffer (50mM NaCl, 50mM TRIS pH 7.5, 1% SDS), then 250 mL of 0.1 mm diameter silica was added. Cells were chilled on ice, then lysed using a bead beater with 3 cycles of 30 sec beating, 30 sec on ice. Lysate was clarified by centrifuging at 21,000 x*g* for 5 min at 4°C and retaining the supernatant.

Peptide Preparation

Cell lysate was quantified using a Pierce BCA assay (Thermo Fisher) per manufacturer's protocols. Lysate was reduced, alkylated, and loaded onto S-Trap mini (ProtiFi) per manufacturer's protocol and digested with 1 µg trypsin in 160 µL of 100mM TEAB pH 8.5 for 2 h at 47°C. Peptides were eluted per manufacturer's protocol and dried down to 20 µL in a vacuum concentrator. *S. typhimurium* peptides were desalted using an Oasis 10 mg HLB desalting column (Waters) and *E. coli* peptides desalted using $C_{18}$ ZipTip (EMD Millipore) per manufacturer's protocol, dried down in a vacuum concentrator, then suspended in 0.1% formic acid for a final concentration of 300 ng/µL.

Serial Dilution of *S. typhimurium*

Desalted *S. typhimurium* peptides were serially diluted two-fold by addition of 60 µL peptides to 60 µL of 0.1% formic acid. Desalted *E. coli* peptides (30 µL) were added to each *S. typhimurium* dilution to create a constant background of *E. coli* peptides, yielding the following fractions of *S. typhimurium* peptides: 0.667, 0.500, 0.333, 0.200, 0.111, 0.059, 0.030, 0.015, 0.008 ,0.004.

Liquid chromatography-mass spectrometry

Technical duplicate injections of 1.33 µL per sample were separated with a PepSep TEN $C_{18}$ 10 cm x 100 µM column (Bruker) and eluted with a 90 min segmented linear gradient from 2-30% ACN. Mass spectra were collected on a Bruker TIMS-TOF Pro operating with the default DDA-PASEF 1.1s cycle time method with two modifications; the CaptiveSpray source set to 1700 V and collision energy maximum to 70 eV.

Identification of mass spectra in FragPipe

Raw data was searched using FragPipe (v.17.1) GUI with MSFragger (v 3.4) and filtered with Philosopher (v 4.2.2-RC).(1, 2) Software parameters for each search are included in the MassIVE repository (See fragpipe.config file). *Escherichia coli* k12 (UP000000625) and *Salmonella typhimurium* LT2 (UP000001014) genomes were downloaded from Uniprot on 2022/03/25. Common contaminants and decoy sequences were added in FragPipe. The two-genome search used default settings, included up to one missed cleavage, oxidation of methionine as a variable modification, no mass-offsets, and was filtered using PeptideProphet with the following command "--nonparam --expectscore --decoyprobs --masswidth 1000.0 --clevel -2 --accmass". The single genome search in MSFragger was set to use the mass-offset algorithm with a corresponding offset for each AAS and the top 26 post-translational modifications discovered in a default setting open search (See Supplementary Table 1). These offsets are available in the SubstitutionOffsets.txt file in MassIVE. The option to "report mass-offset as a variable mod" was set to 1 (Yes – and remove delta mass). Search results were filtered with the same PeptideProphet command. Peptide spectral matches (PSMs) were filtered to 1% false discovery rate (FDR), with no filter for protein FDR. To evaluate decoy PSMs, a separate filtering of the single-genome search used the PeptideProphet command "--nonparam --expectscore --decoyprobs --masswidth 1000.0 --clevel -2 --accmass --minprob 0" and for the Philosopher filter step "--sequential --razor --picked --mapmods --psm 1 --models --prot 1 --ion 1 --pep 1 --protProb 0 --pepProb 0" to remove all PSM filters and recover decoy PSMs.

Genomic analysis to identify single amino acid substitution representing peptides.
To identify a target list of tryptic peptides that differ by one aa between organisms, each genome was digested *in silico* using Protease Guru(3) with the following settings: proteases included trypsin R/KN!P and trypsin R/K, zero or one missed cleavages were allowed, and the minimum peptide length was set to seven. The resultant list of peptides was submitted to a custom Python script (FindSSP.py) that outputs a target list of all peptide sequences that represent a single AAS between the two organisms. The script excludes I/L->L/I substitutions; also R/K->X!R/K at the peptide C-terminus. It results in a comma-separated file of three columns: a redundant list of peptide sequences from the first organism, the corresponding AAS representing sequence from the second organism, and the identified substitution type.

Annotation and filtering of amino acid substitutions
Spectra representing AAS between *E. coli* and *S. typhimurium* were parsed and filtered using a custom Python script (MSFraggerFindSubs.py). To summarize, all PSMs were imported from MSFragger's psm.tsv outputs. Modified sequences were matched from the indicated mass-offset to all considered modifications within 25 ppm peptide mass error. For example, APEPT[-18]IDE would be annotated as 'T->A or Dehydration'. All substituted sequences were then filtered to only include target peptide sequences identified by FindSSP.py. PSMs were filtered to remove 0 intensity spectra, ambiguous or non-AAS modifications.

Annotation of example tandem mass spectrum

The example tandem mass spectrum (see Supplemental Figure 7) was annotated with the Interactive Peptide Spectral Annotator tool. (4)

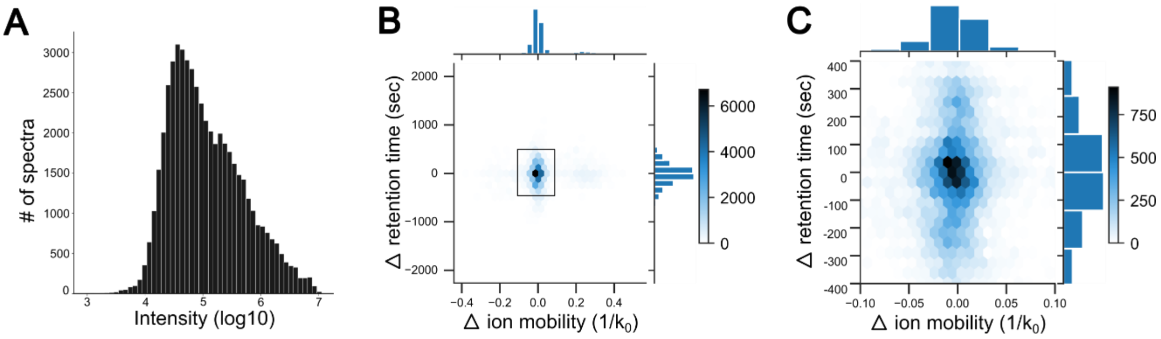


**Supplementary Figure 1. Physiochemical properties of the positive library.** **A)** The intensity distribution of library peptide-spectrum matches. **B)** The shift (Δ) between substitution representing peptide and genomic cognate peptide in retention time (x-axis) and ion mobility (y-axis) for peptides in the positive library with the corresponding distributions in the marginal plots. The box indicates the inset data plotted in **C)**. The color represents the number of peptides in a Δ retention, Δ ion mobility window.

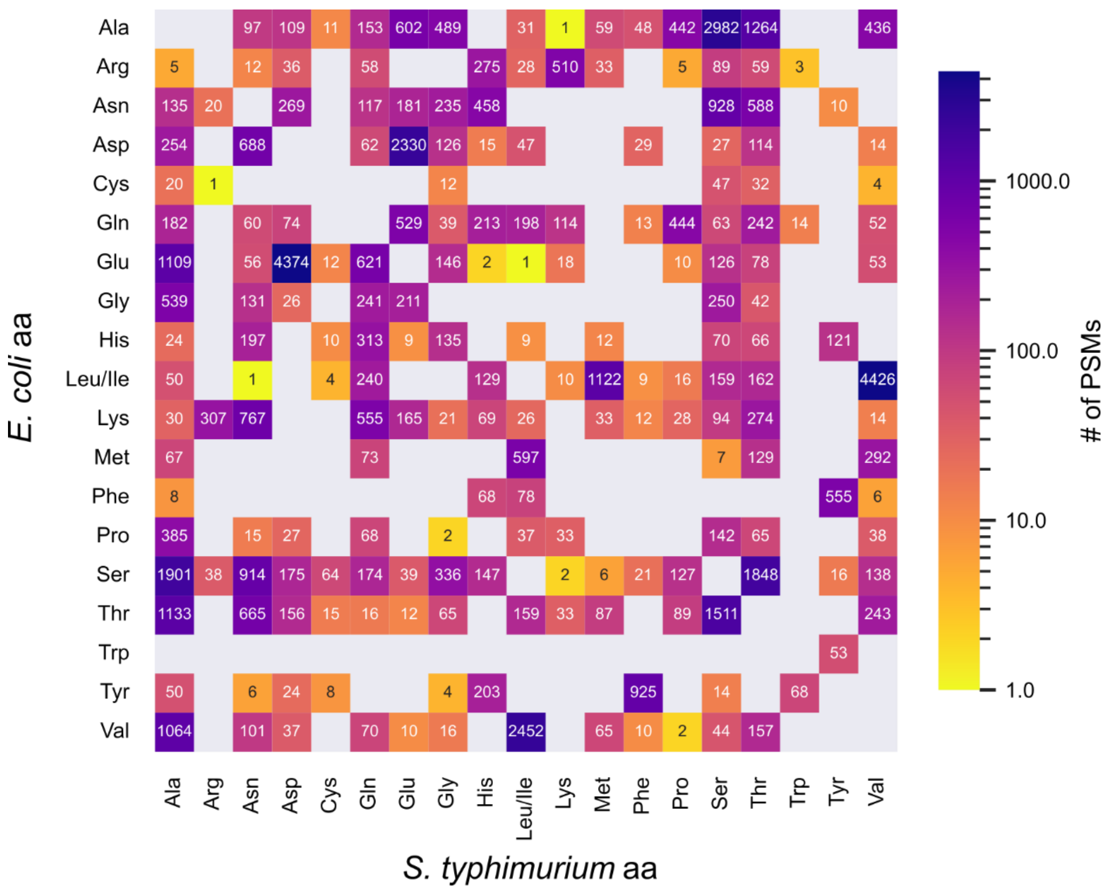


**Supplementary Figure 2. Representation of substitution types in the positive library.** The number of peptide-spectrum matches by substitution type in the positive library, with the genome-anticipated *E. coli* aa on the y-axis and the observed *S. typhimurium* aa on the x-axis. Substitution types not represented by the library are shown in grey.

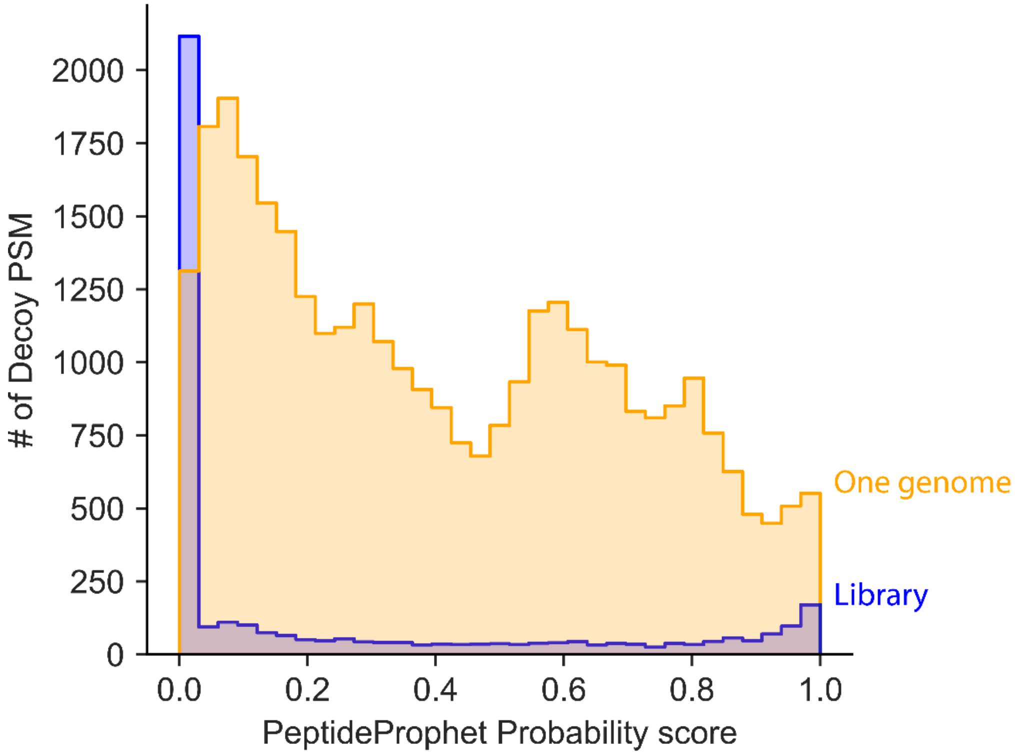


**Supplementary Figure 3. Expansion of the search space increases score threshold for confident sequence assignment.** The PeptideProphet Probability score distribution for decoy peptide-spectrum matches in the library (blue) and one genome (orange) search shows higher scoring decoy hits in the one genome search, suggesting higher score thresholds to maintain a consistent false discovery rate.

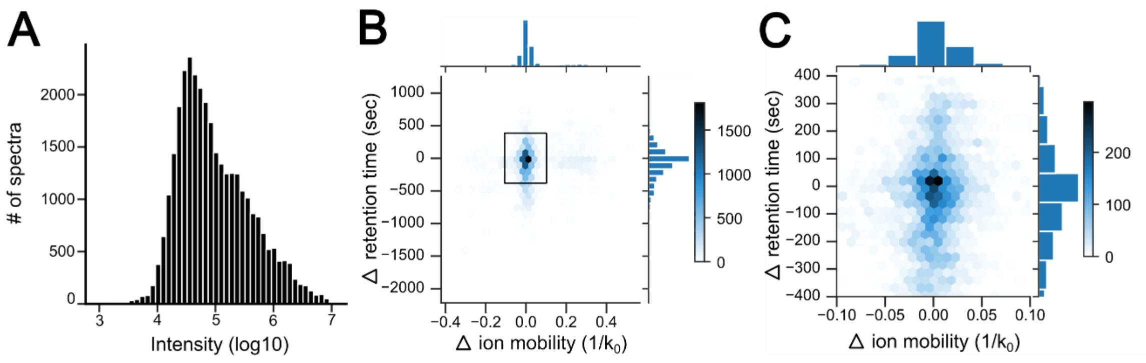


**Supplemental Figure 4. Physiochemical properties of library spectra un- or in-correctly identified in the one genome search. A)** The intensity distribution of peptide-spectrum matches un- or in-correctly identified by the one-genome search. **B)** The shift (Δ) between substitution representing peptide and genomic cognate peptide in retention time (X-axis) and ion mobility (Y-axis) with corresponding distributions in the marginal plots. The box indicates the inset data plotted in **C)**. The color represents the number of peptides in a Δ retention, Δ ion mobility window.

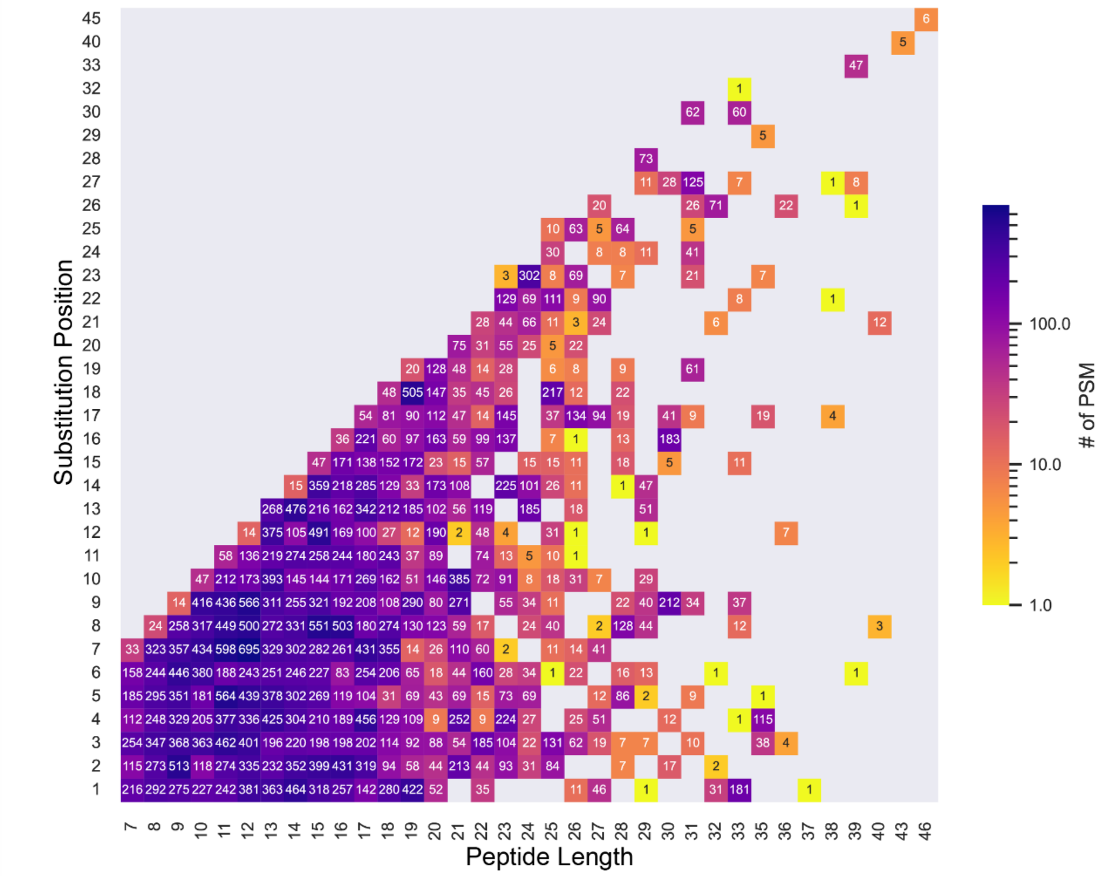


**Supplementary Figure 5. Representation of substitution positions in the positive library.** The number of peptide-spectrum matches in the positive library by substitution position demonstrates representation of common peptide length and substitution position combinations expected in a proteomic experiment. Substitution position and peptide length combinations not represented by the library are shown in grey. Substitution position 1 represents the N-terminal aa; the y=x position represents the C-terminal aa. Note that at the C-terminus we could only identify substitutions that swap lysine and arginine.

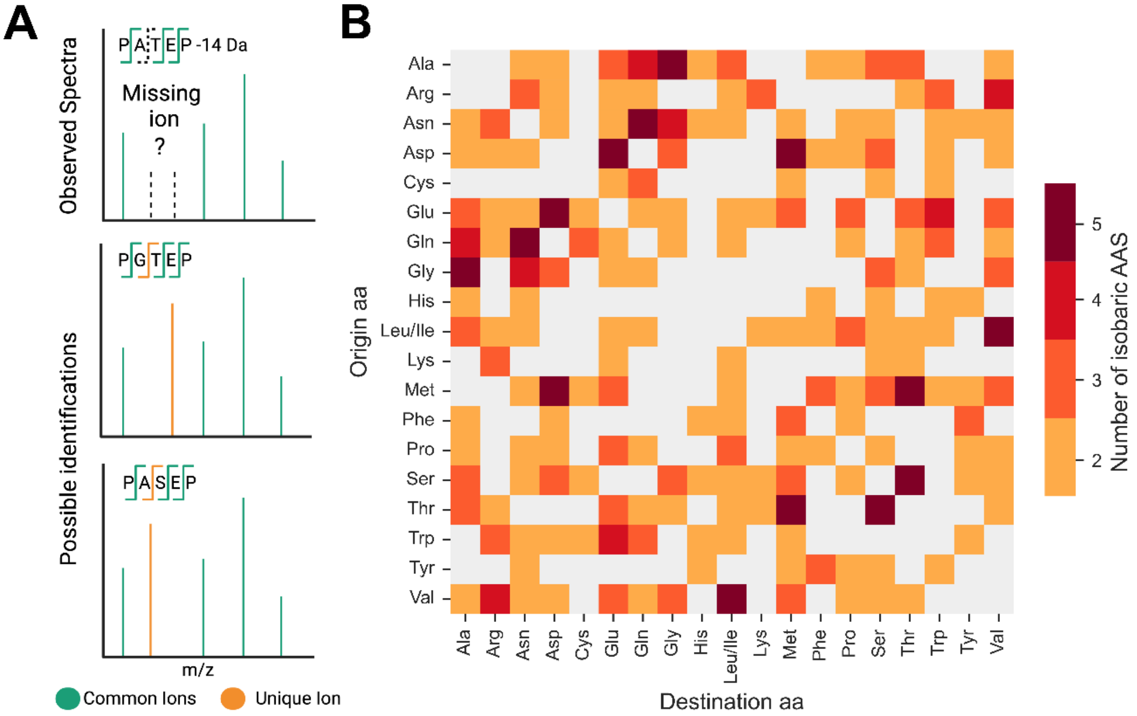


**Supplemental Figure 6. Mass-ambiguity of substitutions. A)** A hypothetical spectrum where unambiguous sequence assignment requires a missing signal unique to a specific fragment (gold) as all other theoretical ions (green) are isobaric between the possible peptide sequences. This occurs with repeated aa, neighboring aa that could have a common PTM of similar mass-offset, and neighboring aa with an isobaric substitution. **B)** The number of isobaric (within 0.02 Da) mass shifts by substitution type. 190/342 AAS types are isobaric with at least one other substitution and may require specific fragment ions for unambiguous sequence determination.

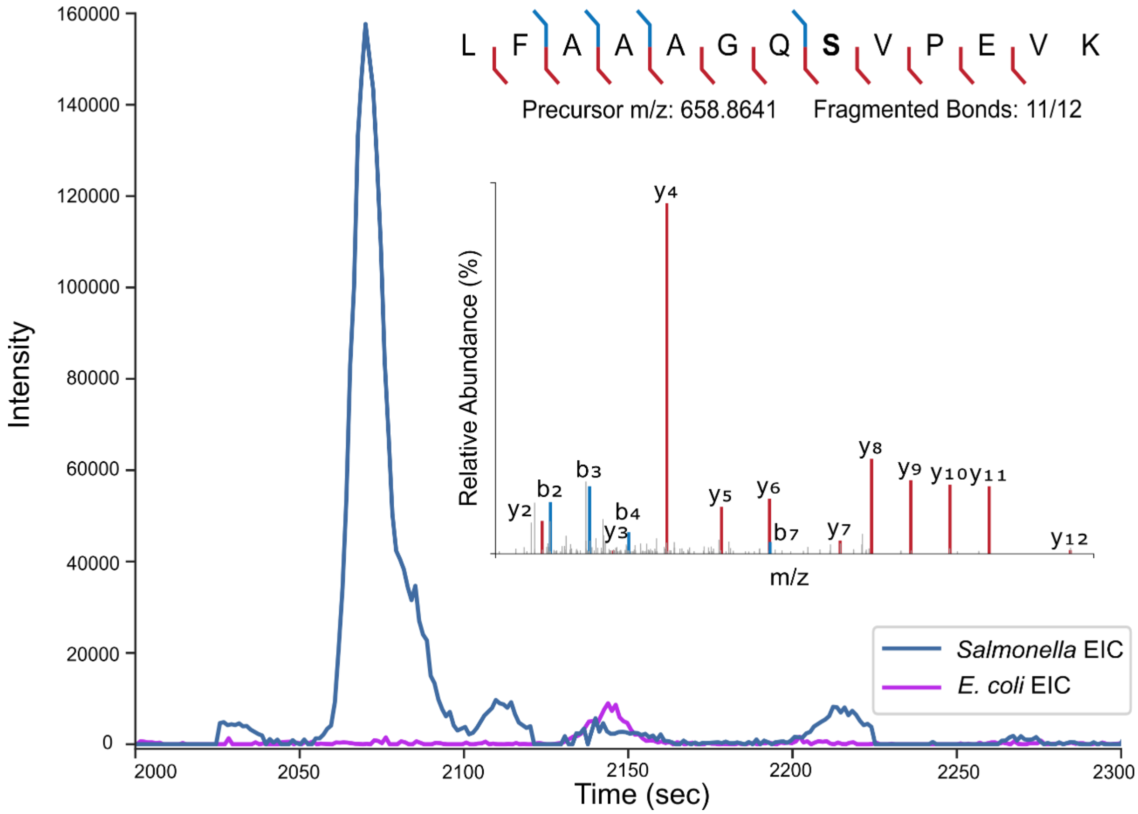


**Supplemental Figure 7. Evidence of a scissor substitution.** The extracted ion chromatogram of 658.86 ± 0.02 m/z is shown in samples containing only *S. typhimurium* peptides (blue) and *E. coli* peptides (purple). Inset is an example MS/MS spectra for the *S. typhimurium* peptide LFAAAGQ**S**VPEVK that exemplifies the loss of a tryptic cutsite in the *E. coli* cognate peptides LFAAAGQK-VPEVK. The bold **S** indicated the amino acid unique to *S. typhimurium*.

Table S1. Common modifications included in the mass-offset list.

| **Modification** | **Mass Shift (Da)** | **Unimod Accession #** |
|---|---|---|
| Failed Carbamidomethylation/Deletion of G | -57.0215 | |
| Homoserine | -29.9928 | 10 |
| Pyro-glu from E/dehydration | -18.0106 | 23,27 |
| Dehydration | -18.0106 | 23 |
| Pyro-glu from Q/Loss of ammonia | -17.0265 | 28,385 |
| Half of a disulfide bridge | -1.00783 | 374 |
| Amidation | -0.98402 | 2 |
| Unmodified | 0 | |
| Deamidation | +0.984016 | 7 |
| First isotopic peak | +1.003355 | |
| Second isotopic peak | +2.00671 | |
| Third isotopic peak/13C3 label for SILAC | +3.010065 | 1296 |
| formaldehyde adduct | +12 | 1009 |
| Methylation | +14.01565 | 34 |
| Oxidation and Hydroxylation | +15.99492 | 35 |
| Sodium adduct | +21.98194 | 30 |
| di-Methylation/Acetaldehyde +28/Ethylation | +28.0313 | 36,255,280 |
| Dihydroxy | +31.98983 | 425 |
| Replacement of proton by potassium | +37.95588 | 530 |
| S-carbamoylmethylcysteine cyclization (N-terminus) | +39.99492 | 26 |
| Acetylation | +42.01057 | 1 |
| Carbamylation | +43.00581 | 5 |
| Replacement of 3 protons by iron | +52.91146 | 1870 |
| Replacement of 2 protons by iron | +53.91929 | 952 |
| Carbamidomethylation | +57.02146 | 4 |
| Addition of lysine due to transpeptidation/Addition of K | +128.095 | 1301 |
| Biotinylation | +226.0776 | 3 |
